# The impact of study design choices on significance and generalizability of canonical correlation analysis in neuroimaging studies

**DOI:** 10.1101/2022.12.14.520275

**Authors:** Grace Pigeau, Manuela Costantino, Gabriel A. Devenyi, Aurelie Bussy, Olivier Parent, M. Mallar Chakravarty

## Abstract

This technical note describes the effects of different data reduction methods and sample sizes for neuroimaging studies in the context of canonical correlation analysis (CCA). CCA is a multivariate statistical technique which has gained increasing popularity in neuroimaging research in recent years. Here, we investigate the parcellation methods’ impact on elucidating neuroanatomical relationships (based on cortical thickness) with known risk factors related to Alzheimer’s disease risk using data from the UK Biobank. The cortical thickness values were parcellated using four common methods in neuroimaging (atlas-based parcellation, spectral clustering, principal component analysis, and independent component analysis) and results from CCA were compared. The results show that the choice of parcellation technique impacts the strength and significance of the correlation between the brain and behaviours. Principal component analysis and independent component analysis result in the strongest correlations.

Additionally, we show that regardless of parcellation technique, smaller sample sizes of participants result in inflated correlation strength and significance.

## Introduction

Canonical correlation analysis (CCA) is a multivariate method used to uncover joint multivariate effects. CCA was first introduced in 1936 (Hotelling, 1936); however, it is more computationally expensive than other standard analysis methods and, as such, has recently become popular due to the availability of increasing computational power. CCA re-expresses data into pairs of high-dimensional linear representations, referred to as canonical variates. Each canonical variate is computed from the weighted sum of the original variable as indicated by the canonical vector.

One of the key advantages of CCA is that it can be used to evaluate two sets of variables without assuming any form of directionality or precedence. In the case of neuroimaging, CCA allows for the simultaneous analysis of a data matrix of brain measurements with respect to a second data matrix of behavioural measures and identifies sources of common variation. CCA then outputs multiple modes (unique pairs of canonical variates) that describe unique patterns of variation in the input sets. The first mode calculated using CCA describes the largest variation in the observed data. The next mode consists of the pair of latent dimensions whose variation between both sets is not accounted for by the first mode, and so on.

CCA produces stable results when the number of observations is greater than the number of features from both input matrices. Previous guidelines have suggested a minimum of between 4 and 70 subjects per feature for CCA (Barcikowski & Stevens, 1975; Leach & Henson, 2014; Thompson, 1990). More recent work suggests that fewer features may be required as the strength of the correlation between the two variable sets increases (Helmer, Warrington, Lisa, Howell, & Rosand, 2020; Yang et al., 2021). The subject-to-feature ratio recommendations are often not fulfilled in neuroscience, where brain voxels are considered individual features. In this case, CCA will still produce results; however, the results are likely to be overfitted to the specific data. As such, data reduction techniques are typically applied before CCA. Principal component analysis (PCA) is the most commonly used data reduction technique and must be applied to the two datasets separately (Zhuang, Yang, & Cordes, 2020). Other techniques include independent component analysis (ICA), feature selection based on statistical dispersion (e.g., mean or median absolute deviation), and the use of brain atlases.

In this study, we use a large subset of data from the UK Biobank (Sudlow et al., 2015) to examine the impact of these data reduction strategies and sample size on the strength of the output correlations, the canonical variates, and the stability of the results. To this end, we examine how these choices impact the results and their interpretation when examining the relationship between neuroanatomy and modifiable risk factors for Alzheimer’s disease (AD). The examination of this relationship is one of the intended uses of the UK Biobank (Sudlow et al., 2015).

## Methods

Canonical correlation analysis (CCA) was run on behavioural and magnetic resonance imaging (MRI) derived metrics from 25,043 participants from the UK Biobank (40–69 years at recruitment) (Sudlow et al., 2015). Behavioural variables chosen correspond with AD risk factors derived from our examination of the literature on modifiable risk factors (Livingston et al., 2020) which were then mapped to the available assessments. Dimensionality reduction was performed on vertex-level cortical thickness (CT) data across participants using four different methods: anatomical parcellation, spectral clustering, principal component analysis (PCA), and independent component analysis (ICA). The reduced CT values were then used as input to the CCA along with the AD-risk factor variables. In the remainder of this technical note, we examine the impact of data reduction techniques on neuroanatomical variables derived from the UK Biobank (UKBB) across a range of sample sizes to determine how these design choices impact the stability and significance of the final results. The methods are described below, and code can be accessed at https://github.com/CoBrALab/repositories/cca.

### Data

The UK Biobank is a representative population-based prospective study with 500,000 participants from the United Kingdom (Sudlow et al., 2015). All participants were aged 40–69 years at recruitment and were first assessed between 2006 and 2010. The assessment visit consisted of physical and cognitive testing, a self-completed touch-screen questionnaire, and a brief computer-assisted interview. A subset of 100,000 participants were selected for follow-up data collection from physical activity monitors and multi-modal imaging, including brain MRI. All released data was collected from three centers dedicated to UKBB imaging using a standardized protocol and data acquired on a Siemens Skyra 3T MR system. The imaging protocol included T1-weighted 3D MPRAGE structural imaging, defaced for subject anonymity and processed before release. The full neuroimaging protocol is provided as part of the UK Biobank Brain Imaging Documentation (Smith, Alfaro-Almagro, & Miller, n.d.). At the time of the study (January 2022), 40,000 subjects had been released and processed using the methods described below.

Epidemiological and cognitive variables collected at the first UKBB assessment were selected to be representative measures of the twelve potentially modifiable AD-related risk factors described in Dementia prevention, intervention, and care: 2020 report of the Lancet Commission (Livingston et al., 2020). These potentially modifiable AD risk factors are low education, hypertension, obesity, hearing loss, traumatic brain injury, alcohol misuse, smoking, depression, physical inactivity, social isolation, diabetes, and air pollution (Livingston et al., 2020). In each of these twelve risk factor categories, different variables were used to describe participant’s health. Variables selected as measures of these risk factors were excluded if more than 10% of the population had missing values. In total, 52 variables were selected and their description are listed in Supplementary Figure 1. Participants who were missing more than 10% (5) of these variables were not included in the study.

The T1-weighted images were processed by CIVET (version 2.1.1; Montreal Neurological Institute; Zijdenbos et al., 2002). CIVET is a fully automated image-processing pipeline for extracting cortical, morphometric, and volumetric data from MR images. Vertex-wise CIVET outputs contain CT estimates at 40,962 vertices for each brain hemisphere (Zijdenbos et al., 2002). Midline non-cortex vertices were removed using a mask, resulting in 38,561 vertices for each hemisphere. Quality control was performed by MC and GP on all the images according to the procedures outlined by the CoBrA Lab to control for accuracy of the surface definition and the GM and WM classifications (see: https://github.com/CoBrALab/documentation/wiki/CIVET-Quality-Control-Guidelines). CT values for all participants were retrieved from McGill University’s NeuroHub portal (https://neurohub.ca/) using the CBrain portal (Sherif et al., 2014). After quality control and removal of participants with missing or unusable data 25,043 participants remained.

### Parcellation Methods

Data decomposition was performed on CT values using four different methods: the Automated Anatomical Labeling (AAL) atlas (Tzourio-Mazoyer et al., 2002), a custom cortical surface atlas grouped via spectral clustering, and two matrix decomposition methods: principal component analysis (PCA) and independent component analysis (ICA). The AAL atlas was selected due to it’s previous use in studies employing CCA, in addition to its popular use in neuroimaging studies in general (Rolls et al., 2020; Zhuang et al., 2020). The parcellation created using spectral clustering was chosen for data decomposition due to previous work highlighting the potential benefits of clustering via spatial proximity (Bhagwat et al., 2019). PCA and ICA were chosen as data decomposition methods due to previous use with CCA for neuroimaging applications (Zhuang et al., 2020). While understanding sex-differences are critical, particularly in the context of AD-related risk, we sought to better understand the stability of the CCA technique based on the design choices that were made. Therefore, the effects of sex were regressed out of the values before all parcellation methods were applied.

The AAL atlas is a popular brain atlas which is widely used in neuroimaging research (Long et al., 2018; Rolls et al., 2020). The AAL atlas is derived from a spatially normalized single-subject high-resolution T1 volume provided by the Montreal Neurological Institute. The AAL template was originally defined on the MNI single brain Colin27 brain (Tzourio-Mazoyer et al., 2002) and registered to the ICBM surface model (Lyttelton et al., 2007). The AAL atlas used for this study partitions the whole cerebral cortex into 90 regions (without cerebellum) which can be seen in Figure 1. For this work, an asymmetric AAL labeling package specific to the resampled surfaces generated from the CIVET pipeline was used. The mean CT of each parcel was calculated for all participants and used as input into CCA in the form of a 90 × 25,043 matrix.

**Figure 1.**
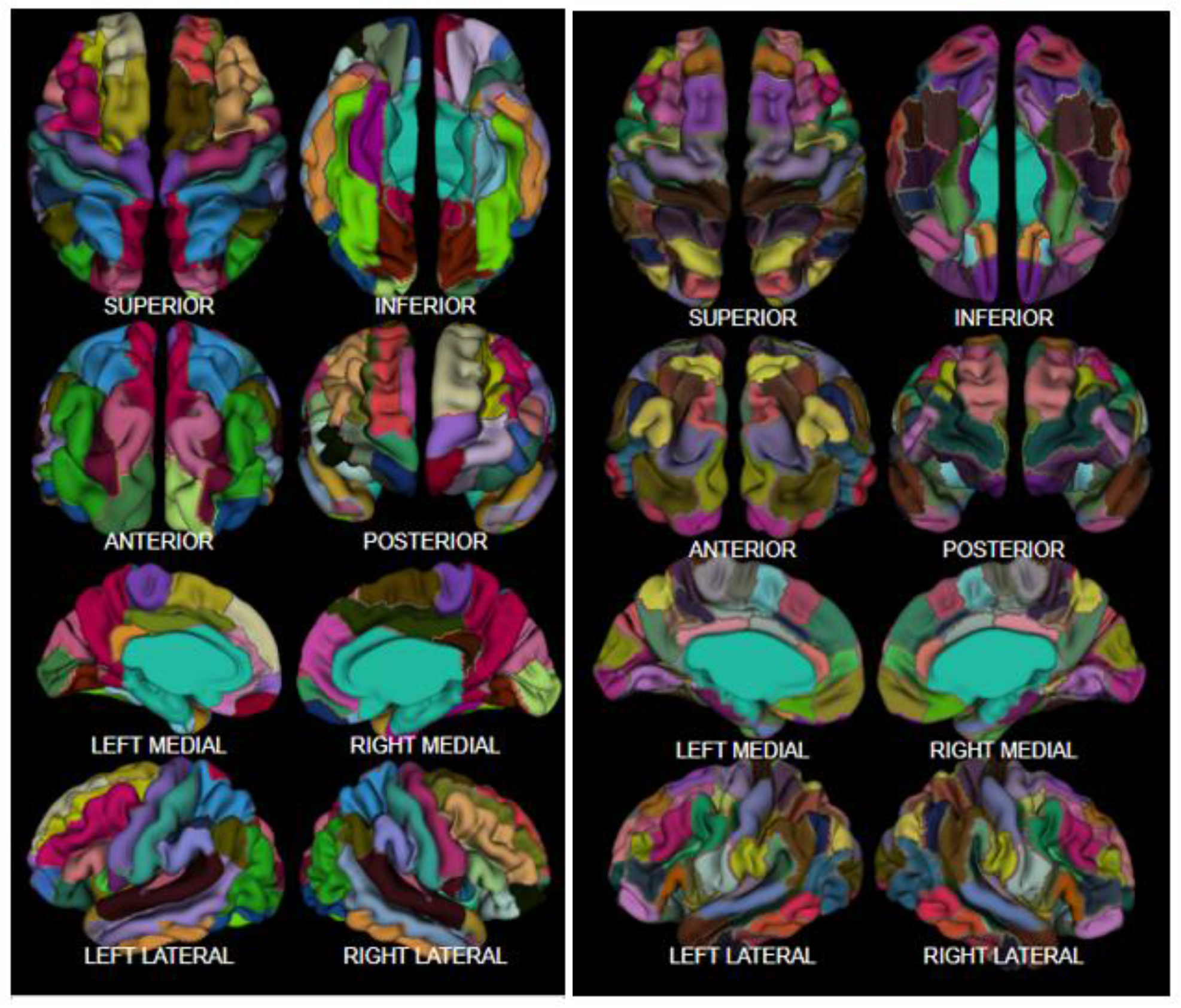
Parcellation Methods. *Left*: AAL atlas parcellations visualized on a brain, showing 90 unique areas. *Right*: Parcellations from spectral clustering visualized on a brain, showing 90 unique areas.

To contrast the anatomical atlas, a spatially derived brain atlas was created using spectral clustering, which does not consider neuroanatomical boundaries; see Figure 1. Spectral clustering allows for the creation of parcels with a similar number of vertices, which allows for unbiased sampling of vertices to estimate CT (Bhagwat et al., 2019). This method uses graph connectivity to identify vertices immediately next to each other which are then mapped to a low-dimensional space that can be easily segregated to form clusters. This was implemented in Python using the built-in spectral clustering algorithm from scikit-learn (version 1.1.1; Pedregosa et al., 2011). A brain mesh is taken as input and each brain vertex is assigned to a discrete parcel. To ensure consistency with the AAL atlas, 90 brain parcels were created and propagated to all participants. The mean CT of each of these parcels was calculated for all participants and used as input into CCA in the form of a 90 × 25,043 matrix. The code used to perform this clustering is available at: _.

Unlike CCA which is scale-invariant, the PCA and ICA matrix decomposition steps performed prior to CCA are influenced by the relative scaling of each variable. Thus, the vertex-wise CT values were normalised using z-scoring by vertex and by subject. PCA performs an orthogonal linear transformation which projects a dataset to a new coordinate system such that the greatest variance of the data is defined on the first coordinate (first principal component), the second greatest variance on the second coordinate, and so on. PCA was run on the vertex-wise CT values for all participants and the top 600 principal components (PCs) were selected as they accounted for over 80% of the explained variance. The participant-specific loadings for all 600 PCs were used as input into CCA in the form of a 600 × 25,043 matrix.

Analogous to PCA, ICA extracts dimensions of hidden variation from high-dimensional variable sets. ICA performs a transformation which projects a dataset to a maximally independent set such that all components are statistically independent and non-gaussian. ICA can be considered as an extension of PCA, however, whereas PCA identifies uncorrelated components, ICA identifies independent components. Additionally, the components of ICA are not naturally ordered. All independent components (ICs) are equally important, thus when ICA was run, 600 components were selected for consistency with PCA. The participant-specific loadings for each ICs were used as input into CCA in the form of a 600 × 25,043 matrix. PCA and ICA were both implemented in Python, the code for which can be accessed at https://github.com/orgs/CoBrALab/repositories/cca/.

### Canonical Correlation

CCA can be expressed as a change of basis. Let ∑_*XY*_be the cross-covariance matrix *cov*(*X, Y*) for any pair of random variables *X* and *Y*. A solution denoting the canonical correlation coefficient *ρ* of the canonical variates can be expressed as follows:

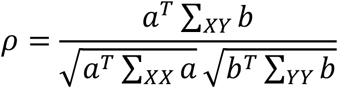

A change of basis can then be defined:

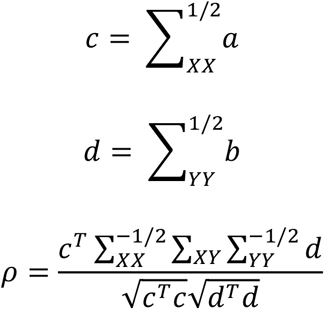

The canonical variates can thus be expressed as:

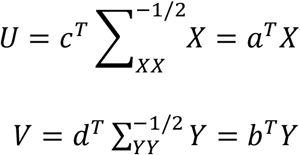

CCA was performed to simultaneously co-analyse each matrix of brain variables along with the matrix of behavioural variables. Each resulting mode from CCA identifies a unique linear combination that relates behavioural variables and CT components. This analysis was implemented in Python using the built-in CCA function from scikit-learn (version 1.1.1; Pedregosa et al., 2011). CCA was used to estimate 52 modes - the minimum size of the two input matrices. The first canonical mode represents the most significant mode of population co-variation, for which individual subjects’ strength of involvement with this mode is highly similar for both a subset of the brain regions and a subset of the behavioural variables. The canonical correlation for each mode (variate pair) was calculated. Canonical correlation between two variates is a primary performance measure for CCA. This measure quantifies the linear correspondence between the two variable sets based on the Pearson’s correlation between their canonical variates.

The traditional approach to interpreting canonical correlation analyses involves examining the magnitude and sign of the canonical weight assigned to each variable in its canonical variate (Lambert & Durand, 1975). However, the use of canonical weights for interpretation or contribution of a variable is subject to criticism (Dattalo, 2014). A small weight may be interpreted as insignificant due its corresponding variable being irrelevant to the overall relationship, or that it has been driven to a small value due to a high degree of multicollinearity with another variable. This same issue is often faced when interpreting the beta weights in regression techniques. Furthermore, canonical weights are subject to considerable instability across samples. These limitations suggest caution in using standardized canonical weights to interpret the results of a canonical analysis (Dattalo, 2014; Lambert & Durand, 1975).

Consequently, canonical loadings have been increasingly used as a basis for interpretation. Canonical loadings measure the linear correlation between an original variable and the canonical variate it belongs to. This measure reflects the variance that the observed variable shares with the canonical variate. In this analysis, canonical loadings were considered to avoid the limitations inherent in canonical weights and to ensure comparisons across different runs of CCA were reliably measured.

After performing CCA using all four methods of data-reduction, the resulting correlations were compared. The Hamming distance between two vectors can be used to compare the order of variables, without accounting for the strength of the loadings. The Hamming distance measures similarity by comparing the changes in the number of positions between the two lists of ordered behavioural variables (Hamming, 1950). Similarly, the normalized Hamming distance is the ratio of the Hamming distance to the length of the lists being compared, with a measure of 0 representing two identical lists. In addition to the normalized Hamming distance, which looks at the relative loadings within the first canonical variates; the correlation between the behavioural loadings themselves can be calculated.

### Permutations and Sampling

Permutation testing using null distributions accounts for the fact that the first modes are expected to have higher correlations (Kernbach et al., 2018; Smith et al., 2015). Here we developed a null distribution against which the real correlations are compared by keeping the behavioural data matrix constant and shuffling the reduced data. For each parcellation method, 5,000 permutations were created and input into CCA. F-approximations of Wilks’ Lambda are used as the test statistic.

To evaluate the effects of participant sample size on the results, CCA was performed on randomly selected subsets of the data. Unlike permutation testing which includes all participants, random sampling includes only a subset of the participants and can be used to demonstrate consistency of results, or a lack thereof, across reduced sample sizes. CCA was run with random samples of 50% (12,521 participants), 25% (6,260 participants), and 10% (2,504 participants) of the data. This was performed using the exact same procedure as running CCA for the full list of participants with all modes of parcellation. The canonical correlation for the first mode was calculated with the reduced sample size. This analysis was also repeated 100 times and averaged for comparison with the first mode from CCA performed on all participants.

## Results

All four parcellation methods resulted in first modes with similar loading patterns. The behavioural variables with the strongest positive and negative loadings are consistent between the four methods; however, the values of the loadings are not. Likewise, similar loading patterns emerge in specific brain areas for all four methods, but the values differ. None of the dimensions of correlation found using any of the parcellation methods were statistically significant. The correlation values for all modes, and the corresponding null mean, are shown in Figure 2.

**Figure 2.**
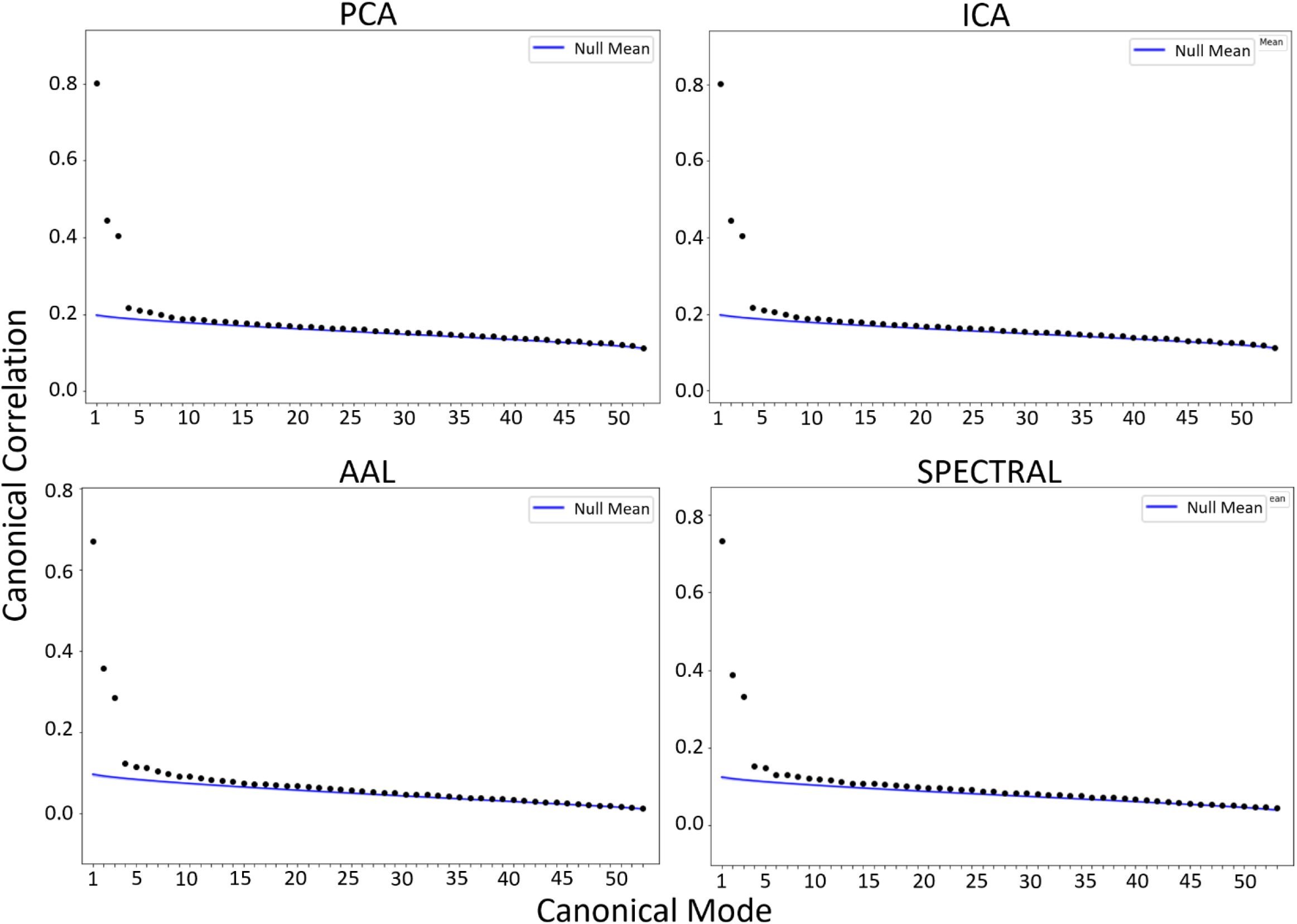
Canonical Correlation of all Modes. The canonical correlation of all modes plotted for each parcellation method used as input into CCA. The blue line shows the mean of the null distribution, estimated via permutation testing.

The behavioural and CT loadings for the first mode found using each method are compared below and can be visualized in Figure 3. The loadings on the behavioural variables in the first mode correlate with the loadings on the CT variables. For example, *low work fulfillment* had the strongest loading across all parcellation methods and was associated with the strong negative loadings aka decreases in cortical thickness. Conversely the negative behavioural loadings can be interpreted as an inverse correlation. For example, high *sport participation* is negatively correlated with CT decreases. This interpretation can be simplified as high *sport participation* is correlated with increased cortical thickness in the specified areas. However, none of the resulting modes from CCA produced significant results and thus making conclusions about these relationships based on the results would not be an accurate representation of the underlying ground truth. CT loadings for individual parcellations can be visualized on a brain map as seen in Figure 3. However, visualizing and interpreting the cumulative effect of the CT loadings on ICs and PCs is intuitively challenging. In addition to the visualization of the strongest loadings in Figure 3, the loadings for all PCs and ICs are shown in Supplementary Figure 2.

**Figure 3.**
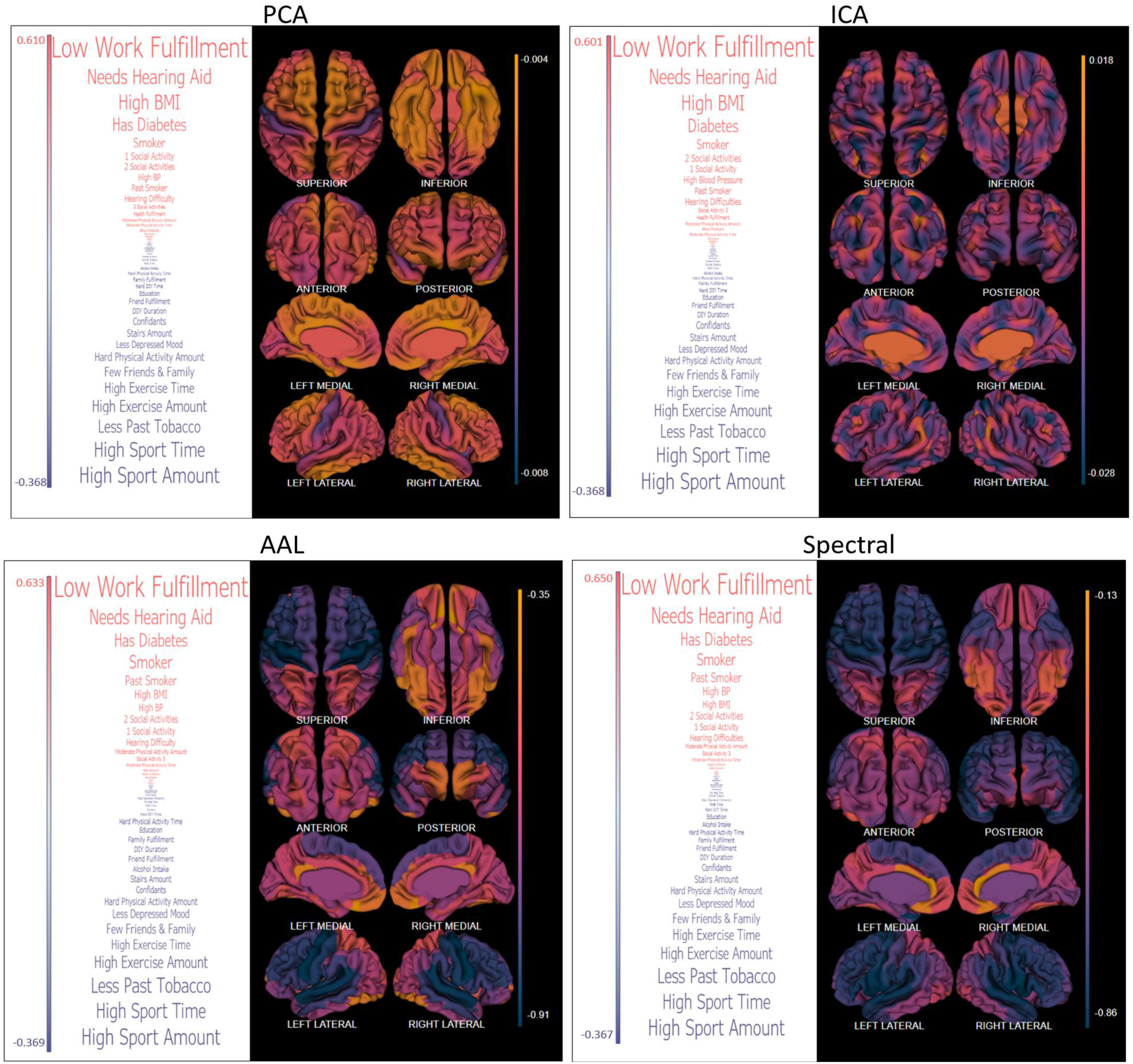
Visualizing Loadings. A visualization of the loadings from the first CCA mode for each parcellation method. *Left*: List of behavioural variables weighted based on their loadings from the first CCA mode. *Right*: Visualization of the spectrally clustered CT value loadings from the first CCA mode. The midline CT values which were not included in the analysis are set to the mean value for

The normalized Hamming distances between the behavioural variables obtained using each parcellation methodology can be seen in Figure 4. The loadings calculated using PCs and ICs are the most similar and both are equally dissimilar from the loadings found using AAL parcellations. Additionally, the distance between the ordered behavioural loadings was calculated for reduced lists. Figure 4 shows the normalized Hamming distance between the top 5 positive loaded variables and bottom 5 negative loaded variables. When considering only the variables with the strongest loadings, the normalized Hamming distance is always lower (the lists are more similar). The lists of behavioural loadings can be seen with their loadings in Figure 4. The matrix of behavioural variables remains the same, only the dimensionality-reduced CT matrices input into CCA are different for each run. Thus, the changes in the behavioural loadings reflect the changes in the relationships uncovered using CCA based on the parcellation method, not any changes in the behavioural variables themselves.

**Figure 4.**
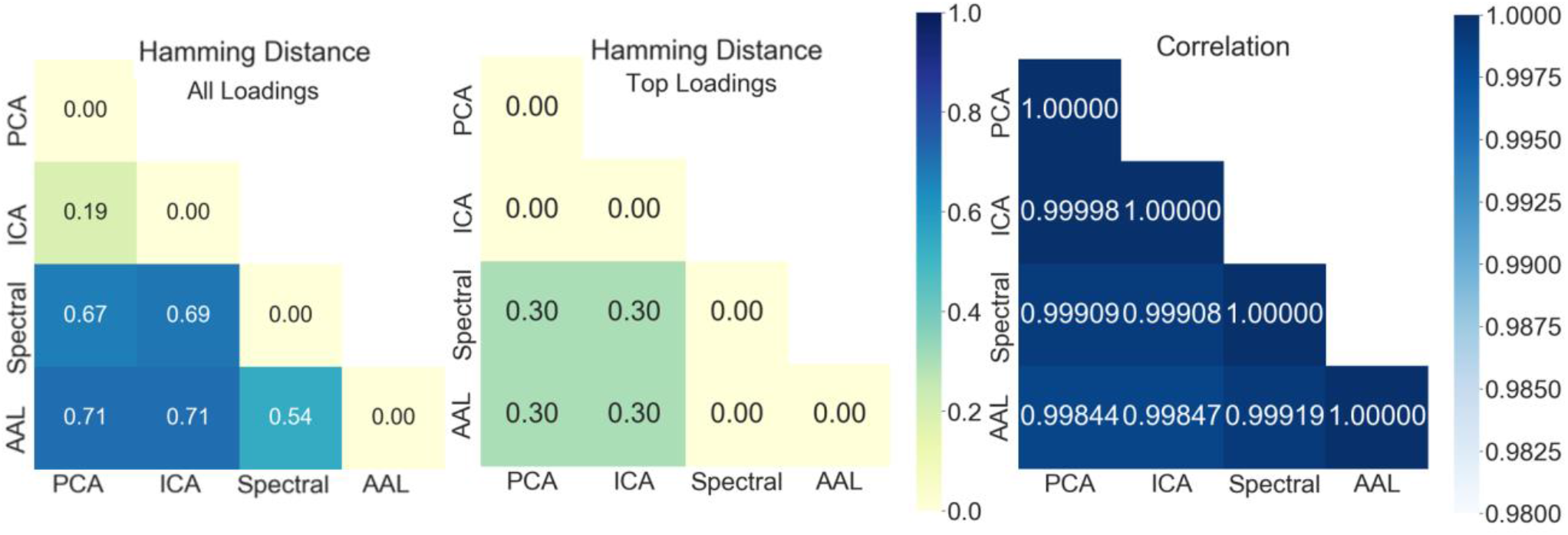
Hamming Distance and Correlation of Loadings. The normalized Hamming distance between all behavioural variables and between the top and bottom five behavioural variables. The correlation between the behavioural variables. Canonical variate loadings are from CCA performed using each parcellation method (specified along the axis) as input.

The correlations between the behavioural variable loadings calculated using CCA with all four parcellation methods can be seen in Figure 4. The correlation values are all high, with the strongest correlations between ICA and PCA and the lowest correlations between ICA and spectral clustering. These correlations measure the strength of the linear relationship between the variate loadings. Large correlations suggest that the loadings are relatively stable across all implementations of CCA (Lambert & Durand, 1975).

The canonical correlation for the first mode of CCA found by averaging the results from 100 random subsamples of the population is visualized in Figure 5. This is shown for the four parcellation methods: PCA, ICA, AAL clustering, and spectral clustering. The corresponding p-values for each sample size are shown in Figure 5 in the upper table. As the sample size decreased the p-value decreased for CCA with all four parcellation methods. The CCA run with PCs an ICs reach significance levels at a sample consisting of 25% of the population.

**Figure 6.**
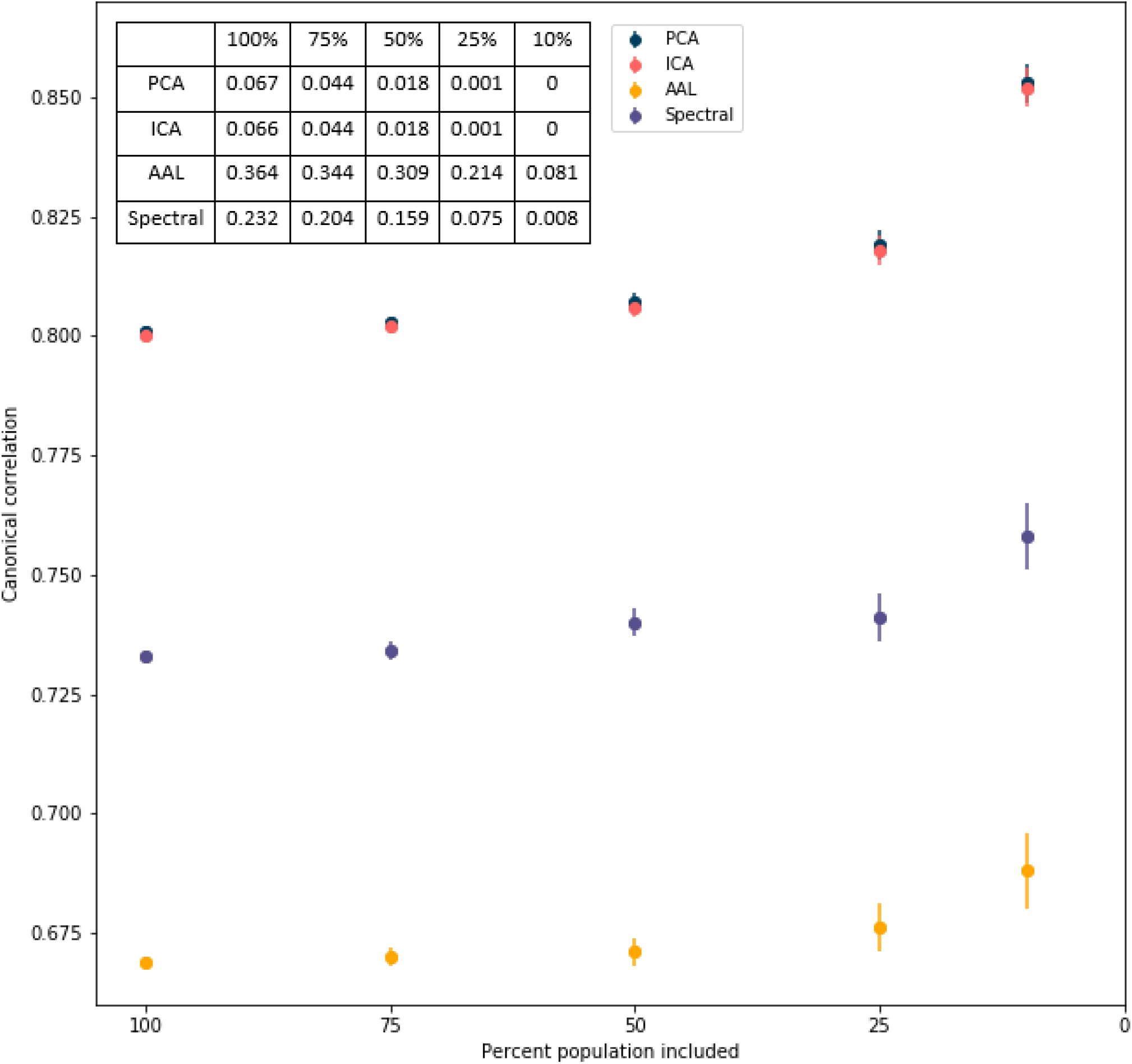
Correlation of Reduced Samples. The average canonical correlation of the first mode found using a subsample of the population. *Top Left*: The average P-values corresponding to the correlations.

## Discussion and Conclusion

None of the parcellation methods resulted in significant CCA results when using the entire sample of participants. As such the relationships between the sets of behavioural variables and brain regions should be interpreted with caution. The resulting modes from PCA and ICA were more significant than those found using the AAL and spectral clustering. This suggests these data-driven, multi-variate data reduction techniques may provide more information and are thus a preferable data reduction technique for use with CCA. The nature of PCA is such that it extracts the most discriminative features, however using PCA can lead to difficulty in interpreting the results. This is likely why CCA papers have significant variation in their choice of parcellation methods (Wang et al., 2020; Zhuang et al., 2020). Additionally, work by Khundrakpam and colleagues suggests that increasing the spatial resolution of a cortical parcellation may improve predictive performance (Khundrakpam et al., 2015). This should be taken into consideration when comparing results found using ICA and PCA (600 components) versus using the AAL atlas and spectral clustering (90 components).

It has previously been suggested that interpretations of a given CCA mode are meaningless if the canonical correlations found are not statistically significant in a smaller validation subsample (Thorndike & Weiss, 1973). Our findings take this a step further and suggest that the canonical correlations of a given CCA mode rely heavily on sample size regardless of the data reduction technique. Our results show that as the size of the random samples of participants decreased, the canonical correlation of the first mode increased. There was also more variation between the samples as the participant size decreased; shown through the increasing standard deviation values. These patterns occurred for all parcellation methods and suggests the canonical correlation of the modes from CCA is not an accurate measure of the correlation between the brain imaging data and behavioural data. This supports similar findings by Yang et al. and has been established theoretically (Q. Yang et al., 2021). The properties of CCA ensure that the first mode found is the maximal correlation of linear combinations of variables between the two input sets. Therefore, the canonical correlation of the first mode approaches one as the number of variables approaches the number of subjects resulting in a full-rank space (Q. Yang et al., 2021).

Although CCA is a promising multivariate approach which holds many advantages in exploring the relationship between the human brain and behavior, it cannot be used or interpreted without restriction. The influence of upstream decisions in the analyses pipeline must be considered when interpreting results and the canonical correlation value should not be interpreted in isolation. All correlations and their reported significance must be considered in the context of dataset dimensionality and subject-to-variable ratio.

