## Supplemental File for "The impact of study design choices on significance and generalizability of canonical correlation analysis in neuroimaging studies"

| CODE | DESCRIPTION | CATEGORY |
|---|---|---|
| 845 | Age completed full time education | Education |
| 864 | Number of days/week walked 10+ minutes | Physical Activity |
| 874 | Duration of walks | Physical Activity |
| 884 | Number of days/week of moderate physical activity 10+ minutes | Physical Activity |
| 894 | Duration of moderate activity | Physical Activity |
| 904 | Number of days/week of vigorous physical activity 10+ minutes | Physical Activity |
| 914 | Duration of vigorous activity | Physical Activity |
| 943 | Frequency of stair climbing in last 4 weeks | Physical Activity |
| 971 | Frequency of walking for pleasure in last 4 weeks | Physical Activity |
| 981 | Duration walking for pleasure | Physical Activity |
| 991 | Frequency of strenuous sports in last 4 weeks | Physical Activity |
| 1001 | Duration of strenuous sports | Physical Activity |
| 1011 | Frequency of light DIY in last 4 weeks | Physical Activity |
| 1021 | Duration of light DIY | Physical Activity |
| 1031 | Frequency of friend/family visits | Social Support |
| 1239 | Current tobacco smoking | Smoking |
| 1249 | Past tobacco smoking | Smoking |
| 1259 | Smoking/smokers in household | Smoking |

| | | |
|---|---|---|
| 1269 | Exposure to tobacco smoke at home | Smoking |
| 1279 | Exposure to tobacco smoke outside home | Smoking |
| 1558 | Alcohol intake frequency. | Alcohol |
| 2050 | Frequency of depressed mood in last 2 weeks | Mental health |
| 2110 | Able to confide | Social Support |
| 2247 | Hearing difficulty/problems | Hearing |
| 2443 | Diabetes diagnosed by doctor | Diabetes |
| 2624 | Frequency of heavy DIY in last 4 weeks | Physical Activity |
| 2634 | Duration of heavy DIY | Physical Activity |
| 2966 | Age high blood pressure diagnosed | Medical Conditions |
| 3456 | Number of cigarettes currently smoked daily (current cigarette smokers) | Smoking |
| 3637 | Frequency of other exercises in last 4 weeks | Physical Activity |
| 3647 | Duration of other exercises | Physical Activity |
| 4537 | Work/job satisfaction | Mental health |
| 4548 | Health satisfaction | Mental health |
| 4559 | Family relationship satisfaction | Mental health |
| 4570 | Friendships satisfaction | Mental health |
| 4792 | Cochlear implant | Hearing |

| | | |
|---|---|---|
| 4803 | Tinnitus | Hearing |
| 6160 | Leisure/social activities | Social Support |
| 20116 | Smoking status | Smoking |
| 20123 | Single episode of probable major depression | Mental Health |
| 20124 | Probable recurrent major depression (moderate) | Mental Health |
| 20125 | Probable recurrent major depression (severe) | Mental Health |
| 20126 | Bipolar and major depression status | Mental Health |
| 23104 | Body mass index (BMI) | BMI |
| 94 | Diastolic blood pressure, manual reading | Blood Pressure |
| 4079 | Diastolic blood pressure, automated reading | Blood Pressure |

**Supplementary Figure 1. UK Biobank Variable Descriptions.** Table containing the codes for each behavioural variable used in the thesis, the UKBB Data Field Description, and the corresponding risk factor associated with the variable.

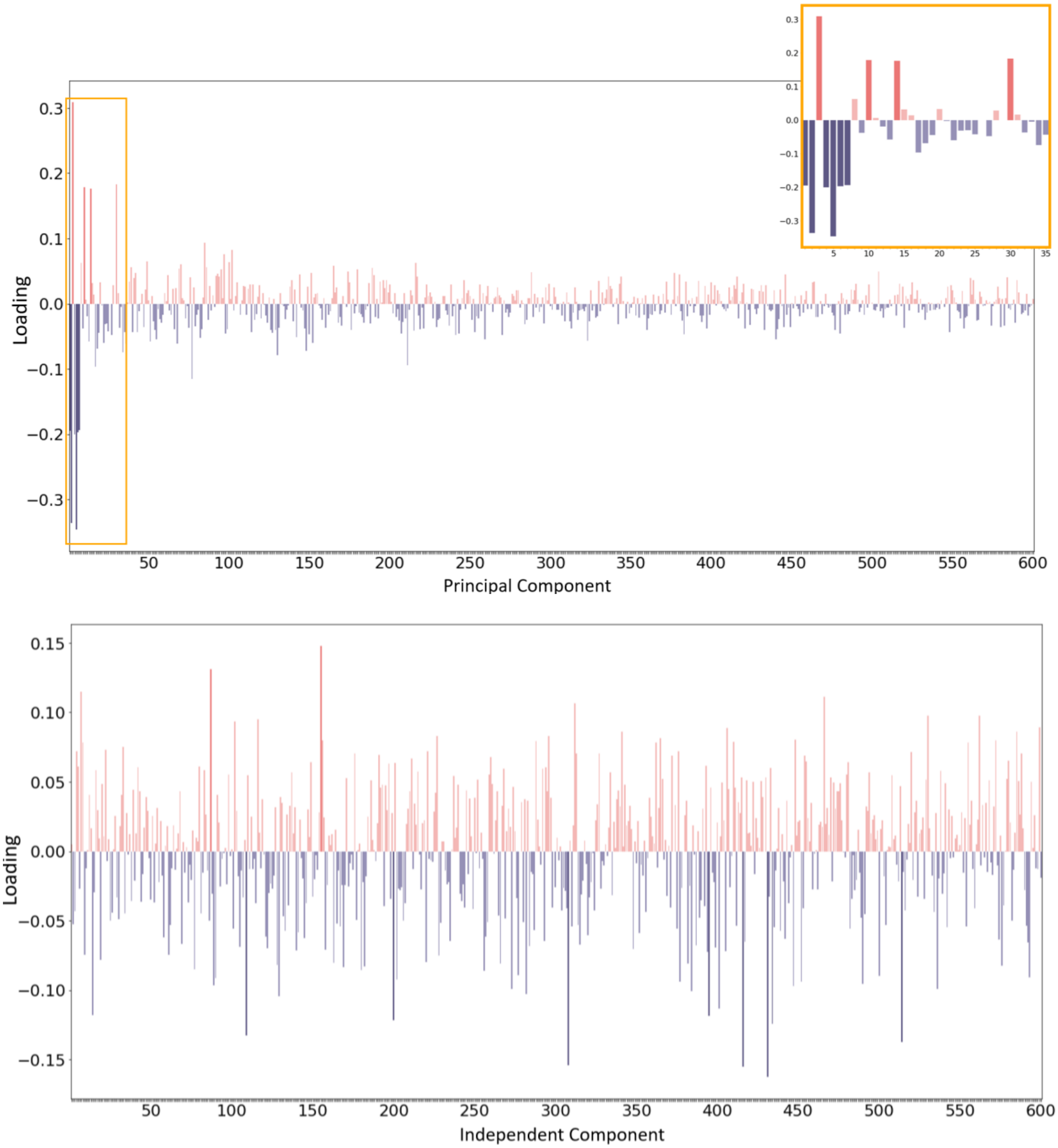


**Supplementary Figure 2. Component Loadings.** *Top*: The loading value for all 600 principal components from the first mode of CCA. The strongest loadings are shown in the top right-hand corner. *Bottom*: The loading value for all 600 independent components from the first mode of CCA.
